# Transcriptomic and functional analysis of *ANGPTL4* overexpression in pancreatic cancer nominates targets that reverse chemoresistance

**DOI:** 10.1101/2022.12.21.521491

**Authors:** Emily R. Gordon, Carter A. Wright, Mikayla James, Sara J. Cooper

**Affiliations:** HudsonAlpha Institute for Biotechnology, 601 Genome Way, Huntsville, AL 35806; The University of Alabama in Huntsville, 301 Sparkman Drive, Huntsville, AL 35899

**Keywords:** ANGPTL4, APOL1, ITGB4, gemcitabine, chemoresistance, pancreatic cancer, PDAC, EMT, transcriptomic analysis, TCGA

## Abstract

**Background:** Pancreatic ductal adenocarcinoma (PDAC) is one of the deadliest cancers based on five-year survival rates. Genes contributing to chemoresistance represent novel therapeutic targets that can improve treatment response. Increased expression of *ANGPTL4* in tumors correlates with poor outcomes in pancreatic cancer.

**Methods:** We used statistical analysis of publicly available gene expression data (TCGA-PAAD) to test whether expression of *ANGPTL4* and its downstream targets, ITGB*4* and *APOL1*, were correlated with patient survival. We measured the impact of *ANGPTL4* overexpression in a common pancreatic cancer cell line, MIA PaCa-2 cells, using CRISPRa for overexpression and DsiRNA for knockdown. We characterized global gene expression changes associated with high levels of *ANGPTL4* and response to gemcitabine treatment using RNA-sequencing. Gemcitabine dose response curves were calculated on modified cell lines by measuring cell viability with CellTiter-Glo (Promega). Impacts on cell migration were measured using a time course scratch assay.

**Results:** We show that *ANGPTL4* overexpression leads to *in vitro* resistance to gemcitabine and reduced survival times in patients. Overexpression of *ANGPTL4* induces transcriptional signatures of tumor invasion and metastasis, proliferation and differentiation, and inhibition of apoptosis. Analyses revealed an overlapping signature of genes associated with both *ANGPTL4* activation and gemcitabine response. Increased expression of the genes in this signature in patient PDAC tissues was significantly associated with shorter patient survival. We identified 42 genes that were both co-regulated with *ANGPTL4* and were responsive to gemcitabine treatment. *ITGB4* and *APOL1* were among these genes. Knockdown of either of these genes in cell lines overexpressing *ANGPTL4* reversed the observed gemcitabine resistance and inhibited cellular migration associated with epithelial to mesenchymal transition (EMT) and *ANGPTL4* overexpression.

**Conclusions:** These data suggest that *ANGPTL4* promotes EMT and regulates the genes *APOL1* and *ITGB4*. Importantly, we show that inhibition of both targets reverses chemoresistance and decreases migratory potential. Our findings have revealed a novel pathway regulating tumor response to treatment and suggest relevant therapeutic targets in pancreatic cancer.

## BACKGROUND

Pancreatic ductal adenocarcinoma (PDAC) is among the deadliest cancers with a five-year survival of only 11% (SEER) [1]. The high mortality of pancreatic cancer patients is explained by the lack of early diagnostic markers leading to late-stage diagnosis of the disease and the high rate of treatment resistance [2]. Standard of care includes resection for about 20% of patients with localized disease who are eligible and treatment with cytotoxic chemotherapies such as gemcitabine or combination agents like FOLFIRINOX (5-FU, oxaliplatin, folinic acid, and irinotecan). This combination, when tolerated, improves progression-free survival by 11 months [3]. Despite the aggressive treatment of this disease with combinations of cytotoxic drugs, the vast majority of patients do not have an enduring response to the treatment. Through improvements in precision cancer treatments based on genetic mutations, implementation of combination therapies like FOLFIRINOX, and advances in immunotherapy over the last decade the five-year survival has increased from 6% to 11%, nevertheless, pancreatic cancer remains extremely deadly. These factors combined with the increased incidence of risk factors like diabetes and obesity. It is predicted that pancreatic cancer will be the second leading cause of cancer deaths by 2030 [4].

Doubling the five-year survival rate is remarkable progress, but novel treatment strategies are still needed. One effective strategy is to identify cellular markers or pathways that are enriched in pancreatic cancer cells. For example, over 90% of PDAC tumors have a mutation in the KRAS gene, however, no KRAS targeting treatments are available and existing targeted therapies like PARP inhibitors [5] and EGFR inhibitors [6] are limited to a subset of patients. It is clear that targeted therapies are impactful for the eligible patients with significant improvement in progression-free survival among patients receiving a targeted treatment [7]. Unfortunately, patients who do not respond to standard chemotherapeutics and are not eligible for existing targeted therapies are left with few options. Thus, there is significant interest in understanding the development of resistance, and in pursuing targets that might sensitize resistant cells to existing treatments.

Our previous transcriptomic study identified genes whose expression was positively or negatively correlated with patient survival and also linked to *in vitro* response to gemcitabine, a common PDAC treatment. We identified ANGPTL4 among those genes whose expression in patient tumors is associated with poor survival and whose knockdown in PDAC cell lines could increase sensitivity to gemcitabine [8]. ANGPTL4 is a member of the family of angiopoietin-like proteins that were first described for their role in angiogenesis [9, 10]. ANGPTL4 can be proteolytically cleaved [11] and the two products each have their own functions. The N-terminal domain plays a role in lipid metabolism and genetic variation in this domain is linked to cardiovascular disease risk. The C-terminal domain has been implicated in tumorigenesis, the promotion of proliferation, and wound healing [11, 12, 13].

Overall, the role of ANGPTL4 in cardiovascular disease and lipid metabolism has been much better described than its roles in cancer. ANGPTL4 has a described role in known cancer pathways including its ability to regulate CREB, FOS, and STAT3 via ERK signaling [14, 15]. ANGPTL4’s ability to alter metabolism and ATP abundance can also impact drug transport [15]. The picture, however, is not perfectly clear since ANGPTL4 expression seems to have different impacts in cancers of different primary sites. For example, methylation and downregulation of ANGPTL4 are associated with progression and metastasis in colon cancer [16] but overexpression is associated with progression and poor prognosis in breast cancer [17]. Breast cancers of the triple negative subtype may be different since ANGPTL4 overexpression in that context has been associated with inhibition of invasion [18]. In pancreatic cancer, *ANGPTL4* overexpression has been associated with tumorigenesis [19], cellular resistance to chemotherapy [8], hypoxia response, and poor patient outcomes [20]. The complicated role for ANGPTL4 motivates our further exploration of its function in pancreatic cancer.

Here, we describe the role of ANGPTL4’s in PDAC by exploring the molecular and cellular changes associated with altered activity of ANGPTL4, the impact of ANGPTL4 on chemoresistance, and the potential for inhibiting downstream pathways driven by ANGPTL4 activation to sensitize tumor cells to treatment. We show that overexpression of *ANGPTL4* leads to chemoresistance, increased migratory potential, and proliferation. Our transcriptomic analysis revealed altered gene expression signatures downstream of *ANGPTL4* overexpression that are linked to epithelial to mesenchymal transition (EMT) and predict patient outcomes. We showed that knockdown of downstream effectors including APOL1 and ITGB4 reversed resistance to treatment and reduced migratory potential. The expression of these genes is also associated with patient survival. These data support the hypothesis that ANGPTL4 and its downstream pathways are potential therapeutic targets for the reversal of treatment resistance in pancreatic cancer.

## METHODS

### Cell Culture

HEK293FT cells (ThermoFisher #70007), MIA PaCa-2 cells (ATCC #CRM-CRL-1420), and Panc-1 (ATCC #CRL-1469) were cultured in D10 media: DMEM (Lonza #12-614Q) supplemented with 10% FBS, and 0.5% Penicillin-Streptomycin. All cell lines were maintained at 37 °C and 5% CO_2_. Cells were cryopreserved with the addition of 10% DMSO (EMD #MX1458-6) to D10 media.

### Plasmids

LentiSAMv2 (Addgene #92062) and lenti-MS2-p65-HSF1-Hygro (Addgene #89308) were used to generate stable cell lines for gene activation. pMD2.G (Addgene #12259) and psPAX2 (Addgene #12260) were used to facilitate viral packaging of sgRNA vector plasmids.

### sgRNA Cloning

sgRNA oligos were designed and cloned into their respective plasmids as described previously (ANGPTL4: 5’-CACGGCCCTGGGGATGCCAAACTGTGG-3’ and NTC: 5’- ACGGAGGCTAAGCGTCGCAA-3’) [21].

### DsiRNA

IDT TriFECTa RNAi kit was used per manufacturer’s protocol. The DsiRNA sequences used were as follows: ANGPTL4 (IDT hs.Ri.ANGPTL4.13.1), APOL1 (IDT hs.Ri.APOL1.13.3), and ITGB4 (IDT hs.Ri.ITGB4.13.3). 100,000 cells were seeded per well of a 12 well tissue culture treated plate 24 hours prior to transfection. Cells were transfected using RNAiMax (ThermoFisher #13778-030) following manufacturer’s recommended protocol. As directed in the TriFecta kit (IDT #hs.Ri.HDAC1.13), TYE 563 transfection efficiency control, positive HPRT-S1 control, and negative (DS NC1) scrambled sequence control were utilized. Functional assays were performed 48 hours after transfection. Expression was validated with each transfection with IDT PrimeTime qPCR Assay system according to manufacturer’s recommendations on an Agilent QuantStudio 6 Flex Real-Time PCR system (ANGPTL4: Hs.PT.58.40146104, ACTB: Hs.PT.39a.22214847, GAPDH: Hs.PT.39a.22214836, and HPRT: Hs.PT.58v.45621572).

### Protein Quantification using AlphaLISA

Briefly, cells were seeded at 25,000 cells in 50uL of media per well of a 96-well tissue-culture treated plate in triplicate per cell type. The following day, the media from each well was transferred to a non-treated 96-well plate and 25ul of AlphaLISA Lysis Buffer (PerkinElmer, #AL001C) was added per well. Plates were then shaken at 250 RPM at room temperature for 10 minutes. Lysates were immediately used. Samples or standards were transferred to a 384 well AlphaPlate (PerkinElmer, #600535, lot:8220-21331) and the AlphaLISA ANGPTL4 manufacturer’s kit protocol (PerkinElmer, #AL3017HV, lot:2905318) was followed for standards, lysate, and supernatant (media) except for the following: lids were covered in foil to prevent light exposure when possible and during incubation periods plates were shaken at 150 RPM for the recommended time. The AlphaLISA plate was read on a BioTek Synergy H5 following previously published protocols [22]. Data were analyzed in R (version 4.1.2) and GraphPad Prism 9.

### RNA-sequencing

Cells were seeded at a density of 0.6x10^6 cells per well of a 6-well plate in triplicate. The following day media was removed and 3ml of media per well with or without 1.5nM of gemcitabine (Sigma #G6423) was added and incubated for 24 hours. Media was removed, cells were washed with PBS (Gibco #10010072), and released from the plate with 2 ml of TrypLE (Gibco #12604-021) per well. After centrifugation, pellets were frozen at -80C until RNA extraction. For RNA extraction, of cell pellets 350 ul of RL Buffer plus 1% BME from the Norgen Total RNA extraction kit and extraction proceeded per manufacturer’s instructions including use of the DNase kit (Norgen # 37500, 25720). RNA quality was verified with the Agilent BioAnalyzer RNA Nano 600 kit (cat# 5067-1512) with the RIN range between 9.2-10. RNA-sequencing libraries were made using Lexogen QuantSeq 3’ mRNA-Seq Library Prep Kit FWD for Illumina kit (cat# 015.24) with 250 ng of RNA input. They were pooled and sequenced on an Illumina NextSeq 500 instrument with 75 bp single-end reads. Read counts averaged 4 million reads and 93.65% of bases exceeding Q30. Lexogen’s BlueBee integrated QuantSeq data analyses pipeline was used for trimming, mapping, and alignment and DESeq2 was used for differential expression [23]. Heatmaps were generated using iDEP.95 [24].

### Scratch-wound Assay

Cells were seeded at 100,000 cells per well of a 96-well plate. After 24 hours uniform scratches were made across the diameter of the wells using a multichannel pipette with 200ul pipette tips and even pressure applied across the wells to cause a wound. The media was then vacuumed off and 200ul of new media was added. Cells were imaged on a Lionheart FX Automated Microscope every 8 hours for 48 hours. Cell culture growth conditions of 37 °C and 5% CO_2_ were maintained throughout. Forty images were taken of each scratch (4 wide by 10 long) with no overlap, autofocus, and auto brightness.

Images were integrated using R and the “tiff” package. Images were further processed using GIMP 2.10. ImageJ and the MRI_wound_Healing_Tool.ijm macro plugin [25] were used to process the images and calculate the area of the wound the cells have not covered. Relative wound closure over time using time 0 for each condition as the control was plotted and a curve line equation was formed by fitting the curve to a non-linear fit one phase decay least squares fit with Yo=0 as a constraint. This base equation (Y=(Y0-Plateau)*exp(-K*X) + Plateau) where Y0 is the Y value when X (time) is zero, Plateau is the Y value at infinite times, and K is the rate constant, was used to determine the value of X or half the time it takes to close the wound. Data were analyzed using GraphPad Prism 9.

### Drug Resistance Screening

Cells were seeded in 96-well plates at 2000 cells/well. Seeded cells were treated with a range of gemcitabine concentrations. Cells were treated again 48 hours later. The number of viable cells surviving drug treatment were measured with CellTiter-Glo (Promega #G7571) 24 hours after the last drug treatment per the manufacturer’s protocol using a BioTek Synergy H5 plate reader. Sample size ranged from 4-11.

### Pathway Analysis

1198 DEG from the ANGPTL4 OE vs KD analysis were imported into the KEGG Mapper-Color [26] and Panther GO Enrichment Analysis [27]. The following parameters were used for KEGG: Search mode: hsa, used uncolored diagrams, included aliases. The following parameters were used for PANTHER: overrepresentation test, homo sapiens reference list, GO biological process complete, Fisher’s exact test, and calculate FDR. Top pathways were merged manually and drawn using BioRender.

### Overall Survival Analysis (OS)

To conduct survival analysis, clinical and RNA-seq expression data was retrieved from The Cancer Genome Atlas for 178 PDAC (TCGA-PAAD) patients (https://portal.gdc.cancer.gov/). Data were normalized using the R package DESeq2 and differentially expressed genes with an FDR < 0.1 were used to generate Kaplan-Meier survival curves. We classified tissues based on their average expression of a given gene set (bottom 25%, middle 50%, and top 25% of gene expression). We compared the patients with the lowest and highest quartile of average gene expression and performed survival analysis. Survival curves and analyses were generated using the “ggplot2”, “survminer”, and “survival” R packages. P values were generated using a log rank test.

### Recurrence Free Survival (RFS)

A Kaplan Meier curve of recurrence-free survival survival plots for ANGPTL4 was created using GEPIA (http://gepia.cancer-pku.cn/) single gene analysis. The relevant parameters were as follows: Disease-free Survival (RFS), Group Cutoff: Quartile (75% high, 25% low), Hazards Ratio: Yes, 95% Confidence Interval: Yes, Axis Units: Months, and datasets Selection: PAAD.

### Correlation Analysis

To compute correlation matrices, RNA-seq expression data was retrieved from The Cancer Genome Atlas for 178 PDAC (TCGA-PAAD) patients (https://portal.gdc.cancer.gov/).

Rank-based correlation coefficients were computed for each of the DEG with log_2_ fold change greater than 0.7 and baseMean>10 (MP2_ANGPTL4_KD vs MP2_ANGPTL4_OE with or without treatment) for 1) all TCGA-PAAD data DEG per gene vs ANGPTL4 expression and 2) all TCGA-PAAD data DEG gene versus OS time. These correlation values were used to generate a list of genes that are co-expressed with ANGPTL4 is overexpressed.

Correlation coefficients and p-values were computed using the "Hmisc" (version 4.6-0) R package. We classified tissues based on their ANGPTL4 expression of a given gene set (bottom 25%, middle 50%, and top 25% of gene expression). We computed correlation matrices using the expression data for patients with the lowest and highest quartile of ANGPTL4 expression which was used to divide samples into quantiles with highest and lowest average gene expression; based on expression of these 42 genes. (40 genes were measured in TCGA: KDM7a and STN1 were omitted).

### Statistical Analysis

Statistical analysis was conducted in R (version 3.6.1 and R version 4.1.2 for RNAseq analysis). The following R packages and software were used for analysis:

survival (version 1.2.1335) [28]

survminer (version 0.4.9) [29]

ggplot2 (version 3.3.6) [30]

DESeq2 (version 1.24.0) [31]

pheatmap (version 1.0.12) [32]

Hmisc" (version 4.6-0) [33]

tiff (version 0.1-11) [34]

## RESULTS

### *ANGPTL4* overexpression increases gemcitabine resistance genes in PDAC

We previously showed that increased *ANGPTL4* expression in tumors is associated with poor survival in PDAC patients (n=51) [8]. Here, we confirm the association with survival in an independent cohort of 178 PDAC patients from The Cancer Genome Atlas (TCGA, PAAD expression dataset). Patients with the highest (top 25%) expression of *ANGPTL4* in tumor tissue had reduced overall survival (OS) and disease-free survival, (p=0.043 and p=0.059 (log-rank test) (**Fig. 1A**, **Supplementary Fig. S1A)**.

**Figure 1:**
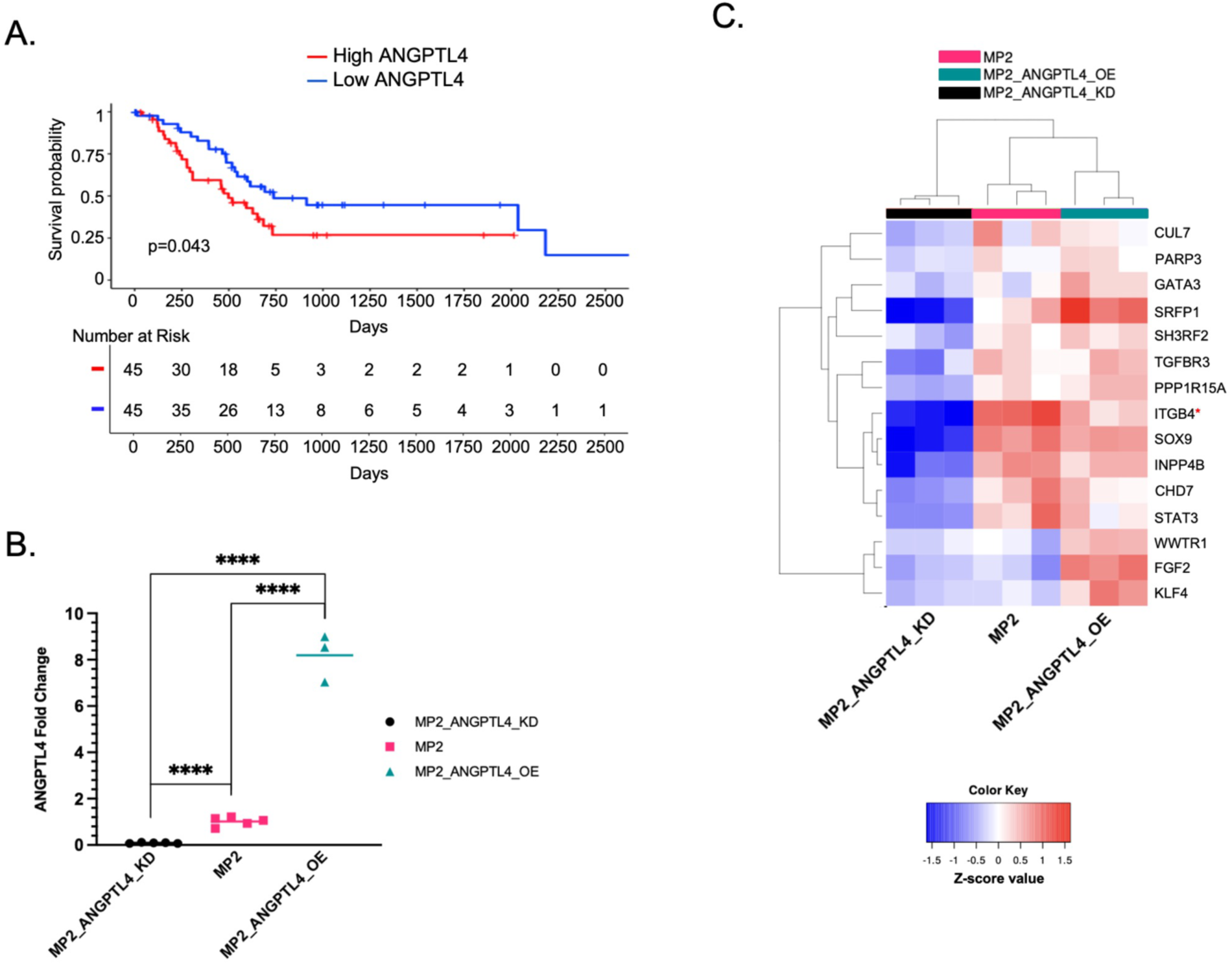
Expression of *ANGPTL4* is associated with patient outcomes and modifies expression of resistance-associated genes. **a)** A Kaplan Meier curve for overall survival using TCGA-PAAD data. The patients with the top (red) and bottom (blue) 25% average *ANGPTL4* gene expression have significantly different OS. (p=0.043, log-rank test) **b)** Relative expression of ANGPTL4 transcript in MP2_ANGPTL4_KD (black circles), MP2 (pink squares), and MP2_ANGPTL4_OE (turquoise triangles) as measured by qPCR. P-values from unpaired, two-tailed parametric t-tests performed with 95% confidence intervals are displayed. Values are normalized with *ACTB* as the housekeeper. (P-values ≤ 0.0001=****) **c)** Heatmap displaying RNA-seq results from MP2_ANGPTL4_KD (black), MP2_NTC (pink), and MP2_ANGPTL4_OE (turquoise) cell lines. Displayed are 15 genes associated with both ANGPTL4 overexpression and gemcitabine resistance. Normalized expression for each of the 15 genes is plotted in the heatmap. Red (*) indicates ITGB4 used for later experiments.

Our previous findings motivated the current study to understand possible mechanisms by which ANGPTL4 contributes to patient survival and specifically how it alters survival outcomes through a possible role in drug resistance. We measured the impact of *ANGPTL4* overexpression and knockdown in the MIA PaCa-2 cell line using CRISPRa (MP2_ANGPTL4_OE) [21] and siRNA knockdown (MP2_ANGPTL4_KD). qPCR showed CRISPRa successfully increased expression of the ANGPTL4 transcript by 8-fold compared to a non-targeting control guide. Similarly, siRNA knockdown reduced expression by over 90% of the control line (**Fig. 1B, Supplementary Table S1**). An ELISA assay showed that protein levels were affected similarly (**Supplementary Fig. S1B**). We measured global gene expression changes using RNA-sequencing on the modified cell lines and controls which revealed 1198 differentially expressed genes (DEG) with a mean read count greater than 10, absolute log_2_Fold Change (log2FC) of at least 0.7, and adjusted p-value (padj) less than 0.05) when comparing overexpressed (OE) ANGPTL4 cells with the knockdown (KD) ANGPTL4 cells (**Supplementary Fig. S1C, Supplementary Table S2**). Given the observed association with drug resistance, we asked whether these DEGs were also associated with gemcitabine resistance by overlapping the 1198 DEG with 114 known gemcitabine resistance genes (**Supplementary Table S3**) [35, 36, 37, 38]. This revealed 15 gemcitabine resistance genes that are altered with overexpression of *ANGPTL4*, 10 of which overlap genes that are part of the EMTome (**Fig. 1C)** [39]. Several of these genes are linked to activation of TGF**β** and its downstream signaling pathways: ERK, PI3K/AKT, and JNK (SH3RF2, TGFBR3, PPP1R15A, INPP4B, WWTR1, and FGF2), contributors to EMT (SOX9, CUL7, and TGFBR3), and transcriptional regulation of pluripotent stem cells (STAT3, KLF4, and FGF2) [26, 27]. Additional pathway analysis showed a connection between the expression of ANGPTL4 and EMT (GO:0001837) in the presence (3.24-fold enrichment, 0.0273 FDR) or absence (3.04-fold enrichment, 0.195 FDR) of gemcitabine treatment (**Supplementary Table S4-S5**).

### Increased *ANGPTL4* expression alters cellular response to gemcitabine

We assessed whether altering *ANGPTL4* expression impacted response to gemcitabine to explore the potential impact of *ANGPTl4* overexpression on treatment response in patients. We measured cell viability in MP2_ANGPTL4_KD and MP2_ANGPTL4_OE cell lines before and after treatment with gemcitabine. As expected, treatment with 12.5nM gemcitabine reduces viability compared to untreated cells. Comparing cells with overexpression of *ANGPTL4* to controls, we observed an increase in cell viability with the same gemcitabine concentrations. Similarly, there was a significant reduction in viability with the *ANGPTL4* knockdown compared to cells overexpressing *ANGPTL4*. This difference in viability is not observed in the untreated cells **(Fig. 2A, Supplementary Table S1)**. These results demonstrate a greater sensitivity to gemcitabine with reduction of ANGPTL4. To understand the link between ANGPTL4 expression and gemcitabine response, we measured transcriptional response to gemcitabine in MP2_ANGPTL4_OE cells and MP2_ANGPTL4_KD cells using RNA-sequencing. We were not surprised to note that 955 genes were differentially expressed upon treatment with the cytotoxic gemcitabine alone. We explored the intersection of the 1198 DEGs identified upon *ANGPTL4* overexpression and the 955 DEGs associated with gemcitabine treatment.

**Figure 2:**
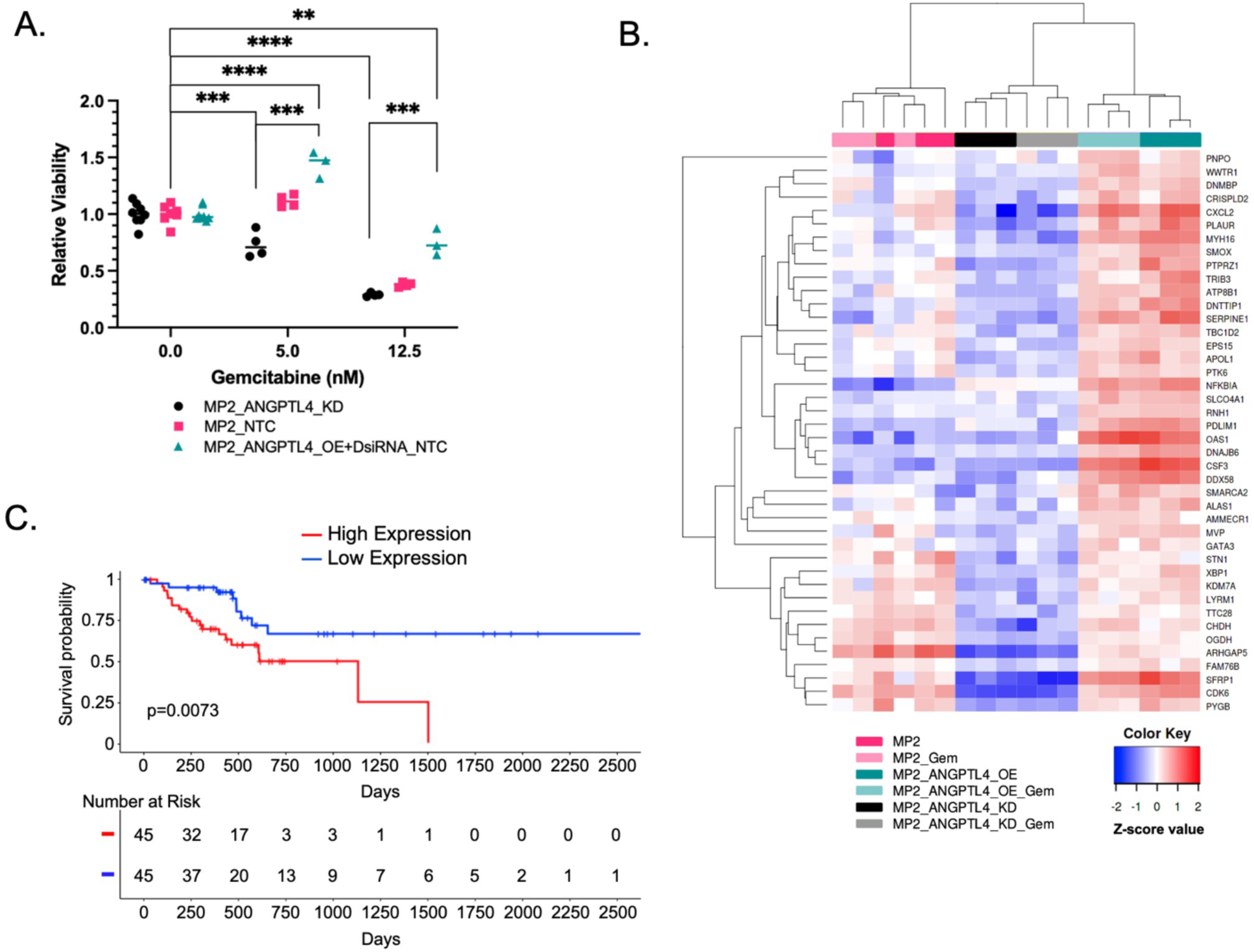
Gemcitabine resistance signature in PDAC cells with high ANGPTL4. **a)** Cell viability was measured for MP2_ANGPTL4_KD (black), MP2_NTC (pink), and MP2_ANGPTL4_OE (turquoise) cells treated with 0, 5nM, or 12.5nM gemcitabine. Relative viability reflects normalization to cells not treated with gemcitabine. P-value asterisks: ≤0.01=**, ≤0.001=*** & ≤0.0001=****, unpaired, two-tailed t-test **b)** Heatmap of gene expression measured by RNA-seq in MP2_ANGPTL4_KD (black/gray), MP2 (pink/light pink), and MP2_ANGPTL4_OE (turquoise/aqua) cell lines both untreated and treated with 1.5nM gemcitabine. Normalized expression of the 42 genes that resulted from an intersection of ANGPTL4 expression DEGs and gemcitabine treatment response DEGs are included in the heatmap. **c)** Kaplan-Meier curve for overall survival using TCGA-PAAD data where patients were divided into quartiles with highest (red) and lowest (blue) using mean expression of the 42 genes in the intersection (40 genes were available in TCGA: *KDM7a* and *STN1* were omitted). P-values were derived from a log-rank test.

This analysis revealed 42 genes associated with gemcitabine response and *ANGPTL4* overexpression (**Fig. 2B, Supplementary Table S6).** This strategy nominated genes that might contribute to altered response to gemcitabine in cells overexpressing *ANGPTL4* and are potentially important for patient outcomes. We tested whether these genes were predictive of patient overall survival using data from TCGA. Since all 42 transcripts were positively correlated with ANGPTL4 expression, we calculated the mean gene expression of these genes and found that patients whose tumors had the highest average expression had significantly reduced overall survival (p=0.0073, **Figure 2C**). The fraction of patients surviving 1500 days or more is dramatically increased in the individuals with low expression of these 42 ANGPTL4 and gemcitabine response genes. This compares favorably to the average 3-year relative survival for PDAC patients in the United States with resectable stage I/II disease-17% [40].

### *ITGB4* and *APOL1* inhibition sensitizes high *ANGPTL4* cells to gemcitabine

I*n vitro* overexpression of *ANGPTL4* alters cellular drug response and correlates with patient survival but the pathway(s) by which this resistance is achieved are unknown. We reasoned that genes co-regulated with ANGPTL4 may function with ANGPTL4 and contribute to resistance and found those genes by identifying genes that are co-expressed with *ANGPTL4* in patient tissues. For each of the 1198 DEGs identified when *ANGPTL4* was overexpressed *in vitro*, we calculated rank-based correlation coefficients and identified co-expressed genes in patient tissues (TCGA-PAAD). For each gene with a significant correlation with ANGPTL4 in patient tissues, we determined whether expression of that gene alone was significantly correlated with patient overall survival (**Supplementary Table S7**). This allowed further filtering of the list based on the assumption that true resistance-associated genes would impact patient survival. From this list, ITGB4 and APOL1 are both positively correlated with *ANGPTL4* expression *in vitro* and in patients, altered with gemcitabine response *in vitro*, and correlated with patient survival in tissues. While there are several genes of interest, we focused on ITGB4 and APOL1 in this study, in part based on their drugability (pharos.nih.gov) and availability of efficient siRNA knockdown reagents. Based on the co-expression with ANGPTL4 and the link to patient survival, we tested whether expression of these genes in combination with ANGPTL4 was relevant for patient outcomes. Comparing tissues that were among the top 25% expression of both *ANGPTL4* and *APOL1* (**Fig. 3A**) or ITGB4 (**Fig. 3B**) to those that were among the bottom 25% expression of both *ANGPTL4* and *APOL1* or *ITGB4* we saw significant differences in overall survival, with overall survival being worse for TCGA-PAAD patients with high ANGPTL4 combined with high APOL1 (p = 0.014) or ITGB4 (p = 0.01) expression. Tumors with high levels (top 25%) of all three genes; *APOL1*, *ITGB4*, and *ANGPTL4,* also had reduced survival compared with the bottom 25%, but when APOL1 and ITGB4 are combined with ANGPTL4 expression the reduction in survival did not become more significant (p = 0.015) (**Supplementary Fig. S1D).**

**Figure 3:**
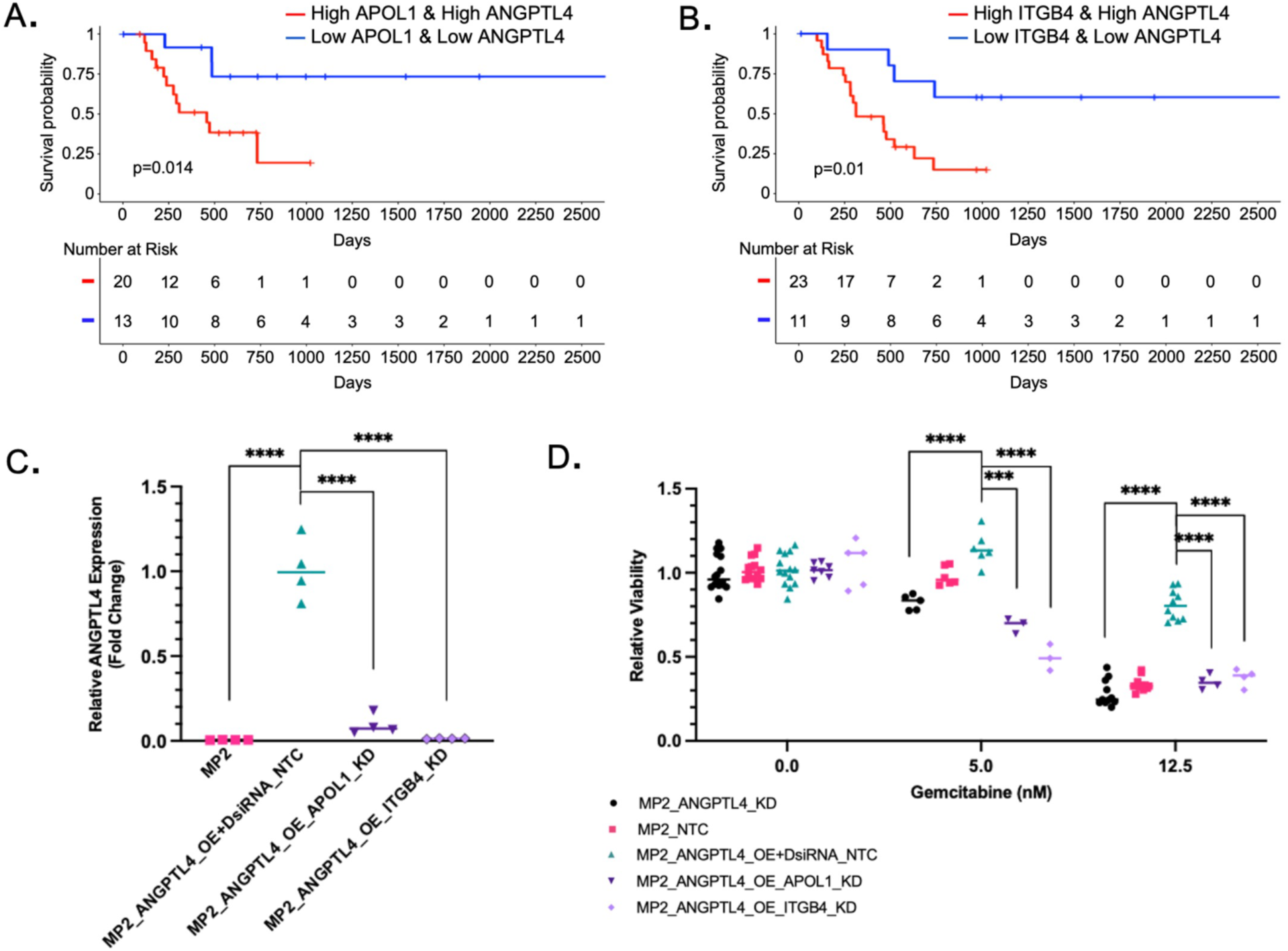
ITGB4 and APOL1 inhibition sensitizes cells overexpressing ANGPTL4 to gemcitabine. **a-b)** Kaplan-Meier plot for overall survival using TCGA-PAAD data. Patients with the top (red) and bottom (blue) 25% average ANGPTL4 gene expression and APOL1 or ITGB4 gene expression (p=0.01 and p=0.014, respectively, log-rank test). **c)** ANGPTL4 expression measured by qPCR normalized to MP2_ANGPTL4_OE+DsiRNA_NTC (turquoise) control: MP2 (pink), MP2_ANGPTL4_OE_ITGB4_KD (lavender), and MP2_ANGPTL4_OE_APOL1_KD (purple) compared to mean of MP2_ANGPTL4_OE+DsiRNA_NTC (MP2_ANGPTL4_OE cells transduced with non-targeting control siRNA). Expression is normalized to the housekeeping gene ACTB. P-values ≤ 0.0001=****; unpaired two-tailed parametric t-test **d)** Relative viability is plotted by normalizing all data to cells not treated with gemcitabine. MP2_ANGPTL4_KD (black), MP2_NTC (pink), MP2_ANGPTL4_OE+DsiRNA_NTC (turquoise), MP2_ANGPTL4_OE_ITGB4_KD (lavender), and MP2_ANGPTL4_OE_APOL1_KD (purple) cells treated with 0, 5nM, or 12.5nM gemcitabine. Sample size ranges from 4-11 per condition. P-value asterisks: ≤0.001=*** & ≤0.0001=****, unpaired, two-tailed parametric t-tests.

Given the association of ANGPTL4 with resistance and patient outcome, we tested whether, like ANGPTL4, loss of *ITGB4* or *APOL1* expression increased sensitivity to gemcitabine *in vitro*. Using siRNAs, we knocked down *APOL1* and *ITGB4* in the MP2_ANGPTL4_OE cell line. We showed that treatment with the siRNAs reduced mRNA levels of the target genes (**Supplementary Figure S2A-B**), and we also found that knockdown of *ITGB4* or *APOL1* reduced expression of *ANGPTL4* significantly compared to a non-targeting siRNA control with ANGPTL4 overexpression background; expression was reduced to 2% in MP2, 5% in ITGB4 knockdown, and 25% in APOL1 knockdown **(Fig. 3C, Supplementary Table S1).** Given our previous finding that overexpression of *ANGPTL4* leads to resistance, we hypothesized that knockdown of *APOL1* and *ITGB4* might reverse resistance given the impact on *ANGPTL4* expression. We tested the impact of *ITGB4* and *APOL1* knockout on gemcitabine sensitivity in the MP2_ANGPTL4_OE line.

After gemcitabine treatment, cells with knockdown of *ITGB4* or *APOL1* in the ANGPTL4_OE background (MP2_ANGPTL4_OE_ITGB4_KD or MP2_ANGPTL4_OE_APOL1_KD) showed reversal of drug resistance when compared to the ANGPTL4 OE cells **(Fig. 3D).** There was no difference in growth rate among the lines that explained the differential viability observed with gemcitabine treatment **(Supplementary Fig. S2C).**

### *ANGPTL4* overexpression increases cell migration

To further understand the pathways that might be involved in resistance, we performed pathway enrichment analysis on the 1198 DEG associated with overexpression of *ANGPTL4* (OE v KD). Those included tumor invasion and metastasis, proliferation, differentiation, inhibition of apoptosis, and cell membrane formation. Genes participating in invasion and metastasis included RhoEGF which was increased with *ANGPTL4* expression. RhoEGF is linked to tumor invasion and metastasis [41]. Many genes in the Ras-ARF6 and Ras-MEK2 signaling pathways are also upregulated (e.g. *FGF, APOL1, ITGB4, EGFR, MPK, CREB, cFOS, JUN*, and *VEGF*) impacting proliferation and differentiation. In addition, ANTPGL4 overexpression is associated with the upregulation of anti-apoptotic pathways including JAK-STAT, Bcl_XS, and SOCS pathways triggered by upregulation of cytokine, cytokine receptors, *CNTF*, and *IL22RA2*. The formation of cellular structure was also affected by *ANGPTL4* and increase in Claudin via the decrease in cell permeability (*GEF-H1, Claudin2*), increased actin assembly (*ARP2/3*), and cell polarity (*AMPK, DLG1*) (**Fig. 4A, Supplementary Table S8).**

**Figure 4.**
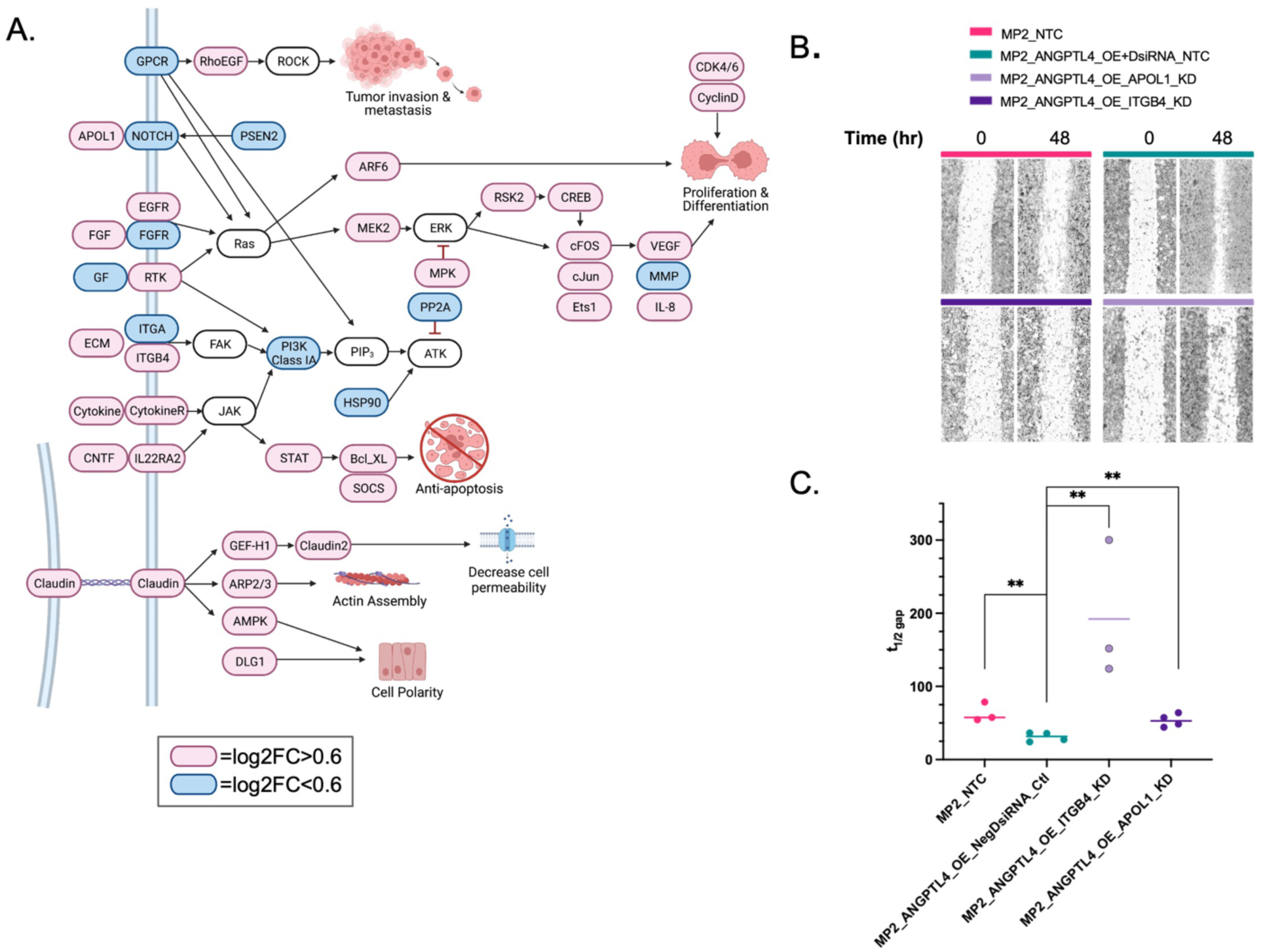
ANGPTL4 overexpression correlation with EMT. **a)** Pathway analysis of 1198 DEG revealed common cancer pathways. Red=log_2_FC >0.7 and Blue=log_2_FC <0.7. **b)** Wound healing assay results at 0 hrs (after initial wound) and 48 hours MP2_NTC (pink), MP2_ANGPTL4_OE+DsiRNA_NTC (turquoise), MP2_ANGPTL4_OE_ITGB4_KD (lavender), and MP2_ANGPTL4_OE_APOL1_KD (purple). Each set was performed in triplicate or quadruplicate. **c)** Time to close half the gap (t1/2 gap) for each of the four cell types is plotted. t1/2: MP2=57.7hrs, MP2_ANGPTL4_OE= 31.6 hrs., MP2_ANGPTL4_OE_ITGB4_KD= 192 hrs., and MP2_ANGPTL4_OE_APOL1_KD=52.96 hrs. P-values ≤0.01=**, unpaired parametric t-test.

Others have shown that on their own *ANGPTL4, APOL1,* and *ITGB4* are each associated with cell proliferation and contribute to PDAC progression [17, 42, 43]. We measured migration using a wound healing time course assay to determine if the increased migration associated with *ANGPTL4* overexpression could be reversed with the KD of *ITGB4* or *APOL1* (**Fig. 4B).** We observed that knockdown of either *APOL1* or *ITGB4* in lines overexpressing *ANGPTL4* reduced the migratory potential of these cells. This was apparent 48 hours after the scratch was made and was quantified by calculating the time to close half of the gap (t1/2 gap) for each of the four cell types is plotted. (**Fig. 4C, Supplementary Table S1).**

## DISCUSSION

Here we have described how overexpression of *ANGPTL4* in pancreatic cancer contributes to disease progression and resistance. We have shown that overexpression of this gene and protein leads to cellular resistance to gemcitabine, one of the most commonly used chemotherapeutics in PDAC. It follows that if overexpression leads to chemoresistance, increased expression of this gene would also be associated with poor patient outcomes, and we confirmed that in independent patient cohorts. The goal of our study is to further understand the role of this gene in chemoresistance in the hopes that a mechanistic understanding will facilitate the development of new treatment strategies. Our transcriptomic analysis revealed that overexpression of *ANGPTL4* broadly impacts transcriptomic profiles in pancreatic cancer cells. This is perhaps not surprising given a rather large body of literature describing not only ANGPTL4’s role in cancer but also cardiovascular disease risk and metabolism [9]. In fact, ANGPTL4’s role in cancer can be difficult to summarize because it has been shown to have both protective and promoting effects in different cancer types [13]. Our initial findings highlighted the potential importance of this gene and existing literature was not sufficient to understand how this gene functions in pancreatic cancer. Using our transcriptomic data, we narrowed our analysis to a list of *ANGPTL4*-impacted genes that are also responsive to gemcitabine treatment *in vitro*. We identified 42 genes associated with *ANGPTL4* overexpression and gemcitabine response and determined that the expression of those genes effectively predicted patient outcomes.

Further narrowing our focus, we identified *APOL1* and *ITGB4* among the genes that are associated with *ANGPTL4* overexpression, gemcitabine resistance, and patient survival and explored them further. We found that knockdown of either *APOL1* or *ITGB4* increased sensitivity to gemcitabine in cells overexpressing *ANGPTL4*. This finding confirms that inhibition of *APOL1* or *ITGB4* can reverse resistance associated with *ANGPTL4* expression in pancreatic cancer cells. One unexpected finding was that knockdown of these genes also reduced expression of *ANGPTL4* itself. Rather than our initial hypothesis that expression of *ANGPTL4* was correlated with *APOL1* or *ITGB4* because they were regulated by *ANGPTL4*, this data supports the hypothesis that *ANGPTL4* is regulated, directly or indirectly, by *APOL1* and *ITGB4* and these genes could be involved in a feedback loop that explains our results.

Considering *APOL1* first, APOL1 (apolipoprotein L1) is part of the HDL lipid complex which plays a key role in lipid metabolism [44]. APOL1 can activate NOTCH1 signaling leading to reduced proliferation and increased apoptosis [42]. NOTCH1 also upregulates ABCC1, a transporter that can also lead to multidrug resistance [45]. In other contexts, NOTCH1 also has been shown to stabilize and activate PPARγ, a transcription factor that promotes transcription of ANGPTL4 [46]. These results suggest a possible mechanism by which knockdown of *APOL1* may reduce *ANGPTL4* expression and further suggests that inhibition of *APOL1*, especially in patients with high expression of *ANGPTL4* might reduce cellular capacity for migration and proliferation. Additional studies would need to be performed to demonstrate whether the link between ANGPTL4 and APOL1 requires NOTCH signaling.

*ITGB4* knockdown also reduces *ANGPTL4* overexpression. ITGB4 (integrin β4, ɑ6β4) is an integrin protein that plays a role in cell-extracellular matrix adhesion. It is upregulated in several cancers including pancreatic cancer and has been linked to poor prognosis, high tumor grade, lymph node metastasis, and drug response [47, 48, 49]. Knockdown of *ITGB4* increases cisplatin sensitivity in lung cancer models [47] suggesting that *ITGB4* is relevant for resistance to multiple chemotherapeutics. ITGB4 may promote receptor tyrosine kinase (RTK) activation through ERBB2 [50]. RTKs generally promote proliferation, invasion, and sustained cell migration, and ITGB4 has a synergistic effect in combination with increased RTKs expression [50, 51]. In our study, we observed increased expression of *CSF3, VEGFA, HBEGF,* and *DDR2*, which all play a role in the epithelial-to-mesenchymal transition in cells overexpressing *ANGPTL4*. An alternate hypothesis is that *ITGB4* promotes cell migration and invasion through regulation of the MEK1-ERK1/2 signaling cascade [49]. Reduction of *ITGB4* in the context of overexpressed *ANGPTL4* could also affect this pathway, explaining the reduced migratory potential of *ITGB4* KD cells in the cells overexpressing *ANGPTL4*. Additional transcriptomic and biochemical studies are needed to determine which of these pathways are most relevant for the chemoresistance phenotypes.

## CONCLUSIONS

Our study provides evidence that *ANGPTL4* overexpression in pancreatic cancer impacts several key signaling and metabolic pathways associated with patient survival, in part through controlling cellular resistance to chemotherapy and cellular migration. *ITGB4* and *APOL1* knockdown are both able to reverse drug resistance and migration increases associated with overexpression of *ANGPTL4* but appear to do so through different mechanisms. These findings emphasize the complexity of developing treatments to target resistance and metastasis; a single approach may not be sufficient. Nonetheless, both *APOL1* and *ITGB4* represent potential novel targets and our data support the further exploration of these genes and the pathways in which they function.

## ABBREVIATIONS

AKT: AKT Serine/Threonine Kinase 1
AMPK: Protein Kinase AMP-Activated Catalytic Subunit Alpha 1
ANGPTL4: Angiopoietin Like 4
ARP2/3: Actin Related Protein 2/3
APOL1: Apolipoprotein L1
Bcl_XS: BCL2 Apoptosis Regulator
cFOS: Fos Proto-Oncogene, AP-1 Transcription Factor Subunit
CNTF: Ciliary Neurotrophic Factor
CO_2_: carbon dioxide
CREB: CAMP Responsive Element Binding Protein
CRISPRa: dCas9 activation system
CSF3: colony-stimulating factor 3
CUL7: Cullin 7
DAVID: Database for Annotation, Visualization, and Integrated Discovery
DEG: differentially expressed genes
DDR2: Discoidin Domain Receptor Tyrosine Kinase 2
DLG1: Discs Large MAGUK Scaffold Protein 1
DsiRNA: Dicer-substrate small interfering RNA
EMT: epithelial to mesenchymal transition
EGFR: Epidermal Growth Factor Receptor
ERK: Mitogen-Activated Protein Kinase 1
ERK1/2: Mitogen-Activated Protein Kinase 1/2
FGF: Fibroblast Growth Factor 1
FGF2: Fibroblast Growth Factor Receptor 2
FOS: Fos Proto-Oncogene, AP-1 Transcription Factor Subunit
GEO: Gene Expression Omnibus
GEPIA: Gene Expression Profiling Interactive Analysis
GEF-H1: Rho/Rac Guanine Nucleotide Exchange Factor 2
HBEGF: Heparin Binding EGF-Like Growth Factor
INPP4B: Inositol Polyphosphate-4-Phosphatase Type II B
ITGB4: Integrin Subunit Beta 4
JAK: Janus Kinase 1
JNK: Mitogen-Activated Protein Kinase 8
JUN: Jun Proto-Oncogene, AP-1 Transcription Factor Subunit
KD: knockdown
KEGG: Kyoto Encyclopedia of Genes and Genomes
KDM7a: Lysine Demethylase 7A
KLF4: KLF Transcription Factor 4
KRAS: KRAS Proto-Oncogene, GTPase
IL22RA2: Interleukin 22 Receptor Subunit Alpha 2
MEK1: Mitogen-Activated Protein Kinase
MP2: MIA PaCa-2 cells
MPK: Mitogen-Activated Protein Kinase
n: number
ng: nanograms
nM: nanomolar
NOTCH: Notch Receptor
OE: overexpression
OS: overall survival
p: p-value
PAAD: pancreatic ductal adenocarcinoma
padj: adjusted p-value
PARP: Poly(ADP-Ribose) Polymerase
PDAC: pancreatic ductal adenocarcinoma
PI3K: Phosphatidylinositol-4,5-Bisphosphate 3-Kinase Catalytic Subunit Delta
PPP1R15A: Protein Phosphatase 1 Regulatory Subunit 15A
qPCR: quantitative polymerase chain reaction
RFS: recurrence-free survival
SEER: Surveillance, Epidemiology, and End Results
SH3RF2: SH3 Domain Containing Ring Finger 2
siRNA: silencing RNA
SOCS: Suppressor of Cytokine Signaling
SOX9: SRY-Box Transcription Factor 9
STAT: Signal Transducer and Activator of Transcription 3
STAT3: Signal Transducer and Activator of Transcription 3
STN1: STN1 subunit of CTS complex
TCGA: The Cancer Genome Atlas
TGFβ: TGFB Induced Factor Homeobox 1
TGFBR3: Transforming Growth Factor Beta Receptor 3
uL: microliter
v: versus
VEGF: Vascular Endothelial Growth Factor A
VEGFA: Vascular Endothelial Growth Factor A
WWTR1: WW Domain Containing Transcription Regulator 1

## DECLARATIONS

### Consent for Publication

NA

### Data Availability

The RNA-seq data generated in this study are available in Gene Expression Omnibus (GEO) at GSE207359. Clinical data and RNA-sequencing data for TCGA-PAAD (PDAC) samples were retrieved on 04/01/2020 using the GDC Data Portal (https://portal.gdc.cancer.gov/projects/TCGA-PAAD). Our analyses included 178 samples in this cohort that had matched clinical and RNA sequencing data.

### Authors’ contributions

ERG, CAW, and SJC designed the experiments. ERG, CAW, and MJ collected data. ERG and CAW analyzed the data. ERG and SJC wrote the first draft. All authors contributed to the writing of the paper and read and approved the final manuscript.

## Acknowledgements

This work was funded by NIH 1R43CA232844-01A1 and the State of Alabama Cancer Research Fund. SJC is supported by UL1TR003096. We would like to thank Anuj Singhal and his team at CFD Research as well as our HudsonAlpha colleagues, especially those in the Myers lab for their feedback, support, and discussions of this work. Figure 4A created with BioRender.com agreement number RD24SQPZ3K.

## Ethics Declaration

All methods were carried out in accordance with relevant guidelines and regulations.

## ADDITIONAL FILES

**Additional file 1: Figure S1.pdf**

**a)** Kaplan-Meier curve of Recurrence/Disease Free Survival (RFS) using GEPIA PPAD dataset. Log rank p=0.047 **b)** Protein abundance of ANGPTL4 in cell lysate and supernatant as measured by AlphaLISA assay for normalization by cell input. MP2_ANGPTL4_OE vs MP2_ANPGTL4_KD p=0.0075 and 0.0018 (**) for lysate and supernatant. MP2_ANGPTL4_OE vs MP2 p=0.0055 (**) for lysate. **c)** Heatmap of RNA-seq data for 1198 DEG from MP2_ANGPTL4_OE vs MP2_ANGPTL4_KD with thresholds: baseMean>10 padj<0.05, and log_2_ fold change +/-0.7.

**Additional file 2: Figure S2.pdf**

**a)** ITGB4 expression measured by qPCR and normalized by the housekeeping gene *ACTB*. All data points are plotted relative to the ANGPTL4_OE line. **b)** *APOL1* expression measured by qPCR and normalized by the housekeeping gene *ACTB*. All data points are plotted relative to the ANGPTL4_OE line. **c)** Cell viability over time normalized to control MP2 cells (lt. pink) at time 0 for MP2_NTC (pink), MP2_ANGPTL4_OE+DsiRNA_NTC (turquoise), MP2_ANGPTL4_OE_ITGB4_KD (lavender), and MP2_ANGPTL4_OE_APOL1_KD (purple).

**Additional file 3: Table S1.txt**

P-values associated with qPCR and viability data

**Additional file 4: Table S2.txt**

DEG from MP2_ANGPTL4_OE vs MP2_ANGPTL4_KD analysis with or without gemcitabine treatment

**Additional file 5: Table S3.txt**

114 Gemcitabine Resistance Genes

**Additional file 6: Table S4.txt**

AMIGO2 Panther GO biological process complete analysis for DEG retrieved from with or without gemcitabine analysis

**Additional file 7: Table S5.txt**

AMIGO2 Panther GO biological process EMT results for with or without gemcitabine analysis

**Additional file 8: Table S6.txt**

List of 42 DEG genes for both ANGPTL4 expression and gemcitabine resistance

**Additional file 9: Table S7.txt**

Correlation analysis of TCGA-PAAD data using DEG gene list from MP2_ANGPTL4_OE vs MP2_ANGPTL4_KD analysis with or without gemcitabine treatment

**Additional file 10: Table S8.txt**

Pathway analysis results from DEG from MP2_ANGPTL4_OE vs MP2_ANGPTL4_KD analysis

---

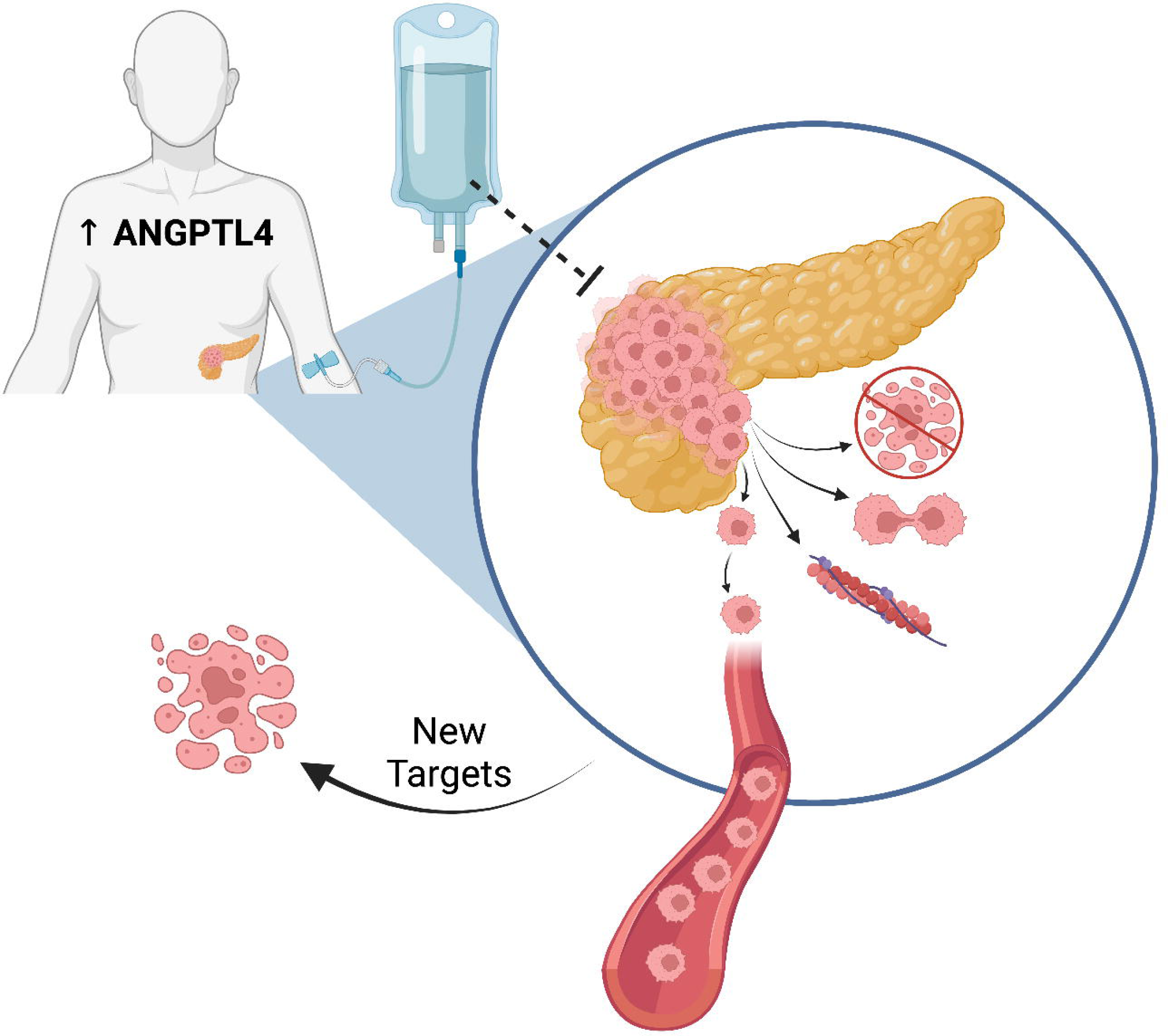

