## Supplemental Figure 2 for "Transcriptomic and functional analysis of *ANGPTL4* overexpression in pancreatic cancer nominates targets that reverse chemoresistance"

**Supplemental Figure 2**: **a)** ITGB4 expression measured by qPCR and normalized by the housekeeping gene *ACTB*. All data points are plotted relative to the ANGPTL4_OE line. **b)** *APOL1* expression measured by qPCR and normalized by the housekeeping gene *ACTB*. All data points are plotted relative to the ANGPTL4_OE line. **c)** Cell viability over time normalized to control MP2 cells (lt. pink) at time 0 for MP2_NTC (pink), MP2_ANGPTL4_OE+DsiRNA_NTC (turquoise), MP2_ANGPTL4_OE_ITGB4_KD (lavender), and MP2_ANGPTL4_OE_APOL1_KD (purple).

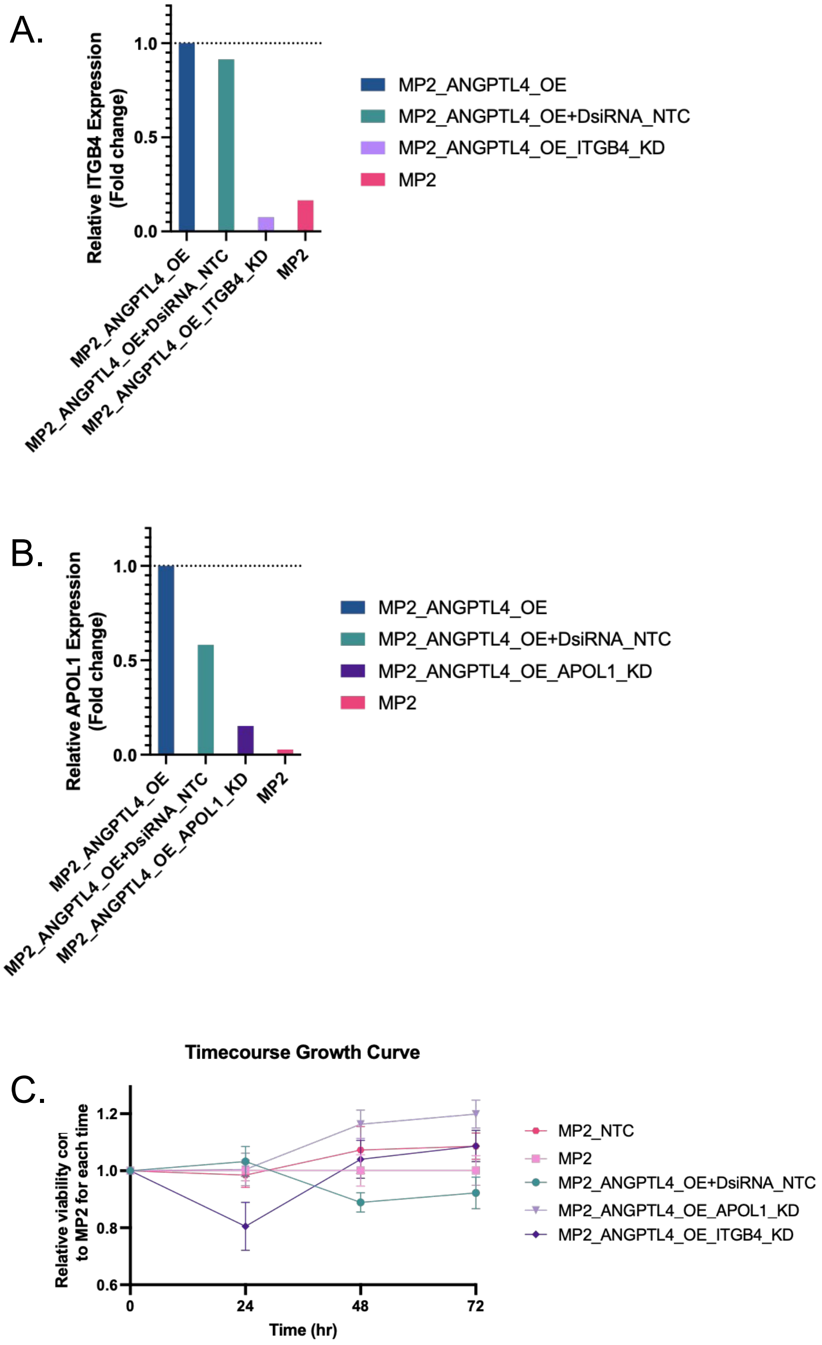
