## Supplemental Figure 1 for "Transcriptomic and functional analysis of *ANGPTL4* overexpression in pancreatic cancer nominates targets that reverse chemoresistance"

**Supplemental Figure S1: a)** Kaplan-Meier curve of Recurrence/Disease Free Survival (RFS) using GEPIA PPAD dataset. Logrank p=0.047 **b)** Protein abundance of ANGPTL4 in cell lysate and supernatant as measured by AlphaLISA assay for normalization by cell input. MP2_ANGPTL4_OE vs MP2_ANPGTL4_KD p=0.0075 and 0.0018 (**) for lysate and supernatant. MP2_ANGPTL4_OE vs MP2 p=0.0055 (**) for lysate. **c)** Heatmap of RNA-seq data for 1198 DEG from MP2_ANGPTL4_OE vs MP2_ANGPTL4_KD with thresholds: baseMean>10 padj<0.05, and $\log_2$ fold change +/-0.7.

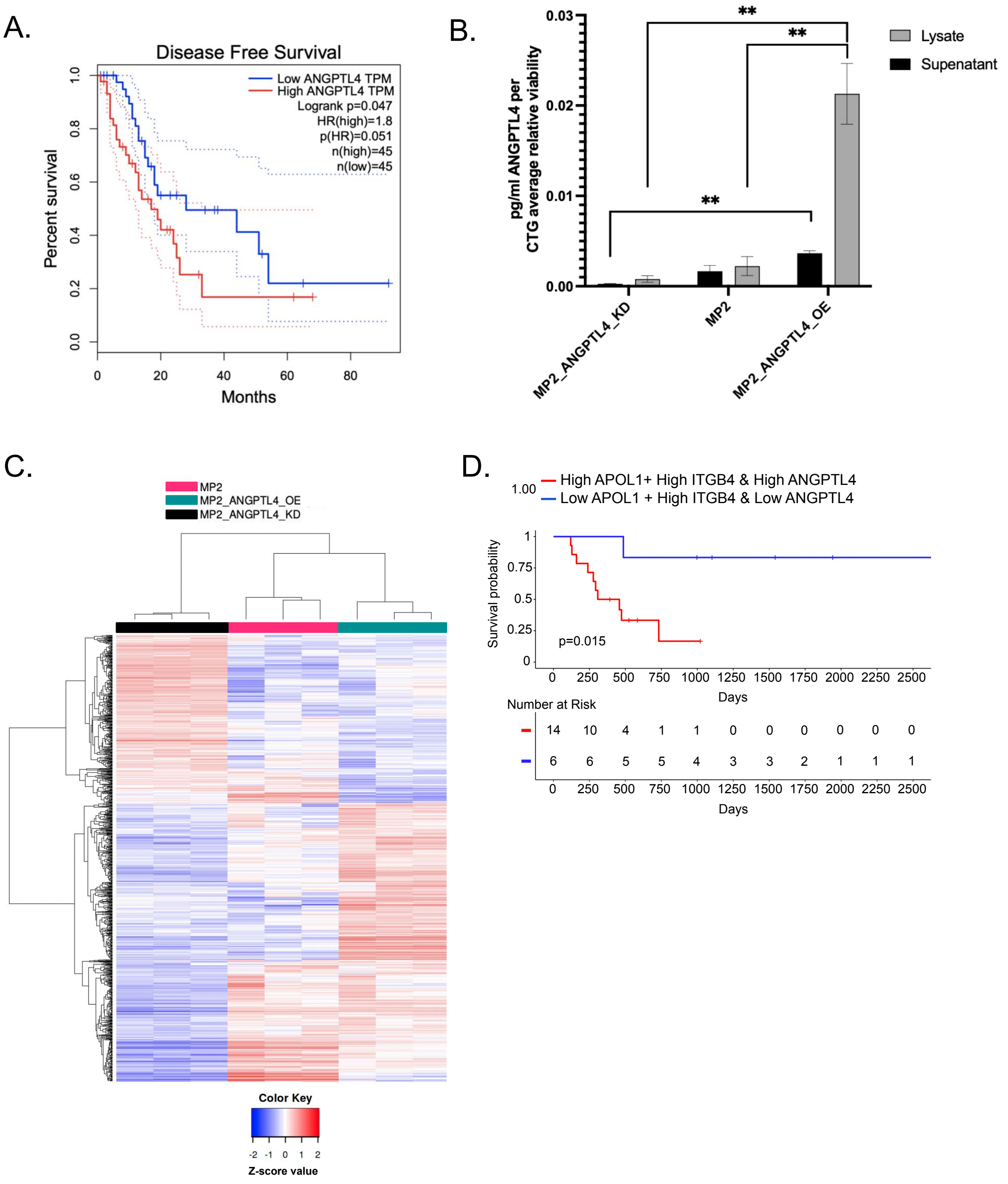
